# Digital-twin optimized transcranial temporal interference electrical stimulation alters rat brain activity patterns and metabolism including deep structures

**DOI:** 10.64898/2026.09.26.754615

**Authors:** Patricia Wenk, Keivan Ahmadi, Daniel Mertens, Fahmida Akter, Aref Kalantari, Annika Michalek, Tino Zaehle, Markus Aswendt, Philipp Ruhnau, Rolf Findeisen, Eike Budinger

## Abstract

Non-invasive deep-brain stimulation (DBS) using transcranial alternating current stimulation (tACS) with temporal interference (TI) has emerged as a promising approach for therapeutic applications in motor and neuropsychiatric disorders. Here, we present a preclinical TI tACS protocol for adult rats that follows a systems approach: an anatomically realistic finite-element digital twin of the rat head is used to optimize electrode placement and injected currents for a deep target, and the model predictions are then tested *in vivo*. We combine, minimal invasive TI tACS with functional MRI (fMRI) to characterize stimulation-related brain-wide neural responses, alterations in functional connectivity, and changes in neurotransmitter levels. The optimized montage targeted in particular the ventral (motor-related) thalamus. Stimulation-evoked fMRI revealed robust bilateral activation across both subcortical and cortical regions, such as the thalamus, hippocampus, amygdala, retrosplenial and anterior cingulate cortex, which, in turn, reflects direct and indirect activations of brain circuitries by TI tACS. Changes in the connectivity patterns during resting-state fMRI were limited (e.g., fimbria of the hippocampus), but partially overlapped with regional stimulation-evoked fMRI. Therefore, we assume only a short-term influence of TI on neural networks following a single stimulation. MR spectroscopy indicated a trend towards an increased glutamate/GABA ratio in the thalamus. Overall, we demonstrate that TI is a feasible and effective approach for non-invasive DBS in rodents, and that digital twin-based optimization combined with *in vivo* validation provides a workflow towards individualized stimulation and a foundation for future mechanistic and translational studies in disease models.

**Highlights:**

- Digital twin modeling optimizes TI tACS for deep-brain stimulation in rats
- TI tACS evokes robust bilateral cortical and subcortical fMRI responses
- TI tACS induces direct and indirect activation of distributed brain circuits
- Resting-state connectivity changes after a single TI session are limited
- TI tACS tends to increase the thalamic glutamate/GABA ratio

**Graphical Abstract:** 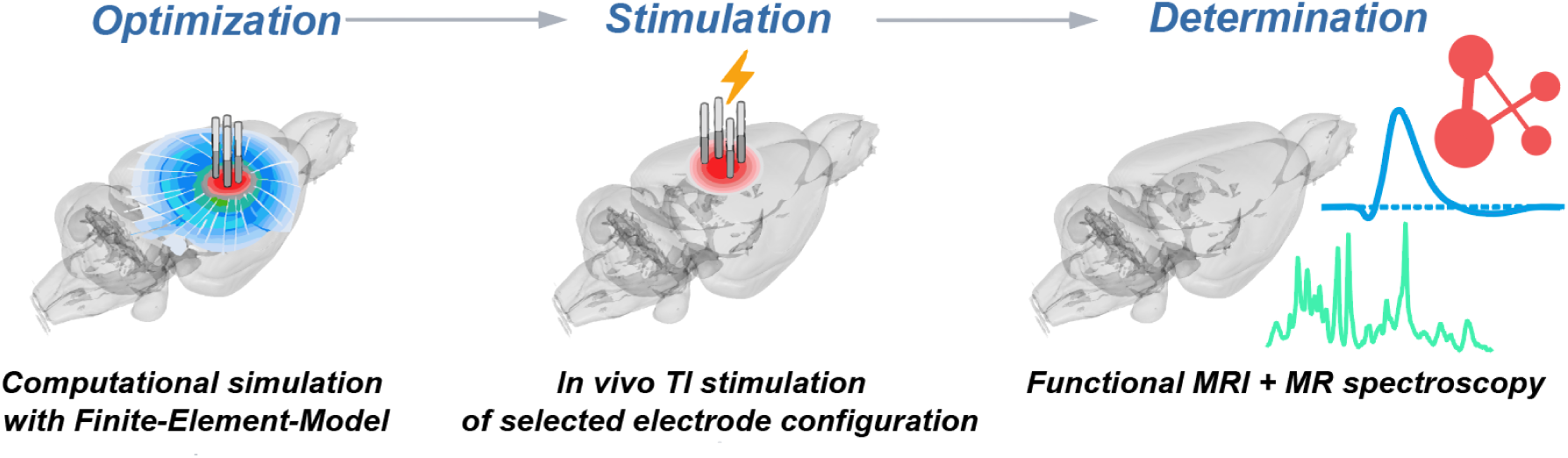

## Introduction

Deep brain stimulation (DBS) is a common therapeutic approach to target symptoms of motor disorders such as essential tremor [1], dystonia [2], and Parkinson’s disease [3]. In a recent study, Hacker and colleagues reported a long-term benefit when DBS was applied to early-stage Parkinson’s disease compared to conventional medication [4]. However, state-of-the-art DBS is based on an invasive surgical procedure potentially leading to severe complications [5]. To reduce such risks and improve the patients’ well-being, it is necessary to investigate possible adaptations to this procedure.

One approach is to use non-invasive electrical stimulation, for instance, transcranial direct current stimulation (tDCS) or transcranial alternating current stimulation (tACS). tDCS was already successfully applied to boost temporary learning acceleration and memory performance [6], and to reduce symptoms of depression [7]. However, tDCS primarily induces polarity-dependent changes in cortical excitability within targeted regions. While different electrode montages, including bilateral configurations, are possible, its effects are generally not frequency-specific. In contrast, tACS applies oscillatory electric fields that can entrain neural activity, enabling the modulation of brain oscillations within and across regions [8]. This effect makes tACS suitable to reset disturbances in brain oscillations and to treat neuropsychiatric disorders, such as schizophrenia [9,10]. Though, tACS is limited to cortical regions and not suitable for deeper brain structures. Recently, Grossman and colleagues combined tACS with temporal interference (TI) to successfully stimulate hippocampal neurons in mice [11]. Here, two pairs of scalp electrodes with alternating sinusoidal waves (typically in the kHz range) with a small difference in frequency (e.g., 10 Hz) lead to a superposition of the electrical fields and enhancement of the interfering currents in a target area. TI has become an emerging technique in the field of non-invasive DBS. Several computational modelling approaches validated this concept, e.g., in murine and human head models, where a manipulation of the strength of the applied electrical fields can reach large interference in deep brain structures like the thalamus with little interference in more superficial areas [12], and model-based optimization of TI electrode montages has been proposed to make such targeting systematic [13]. Importantly, Alekseichuk and colleagues reported an up to 100-fold decrease in electrical field strength with increasing head size from mice to monkey and human heads. This indicates a strong specimen-dependent outcome and therefore the need to develop an individual experimental setup for different species [14] or individual subjects [15].

In very recent studies, modified versions of the original TI protocol have been successfully applied to improve spatial navigation behavior in humans [16] or motor performance in mice [17]. Combinations of TI with functional magnetic resonance (fMRI) and behavior demonstrated (I) a hippocampal stimulation, which was in line with an increased performance in an episodic memory test in healthy subjects [18], and (II) a striatal stimulation related to enhance performance in a motor task in healthy older participants [19]. Recently, TI-induced amplitude modulation, neuronal entrainment, and hemodynamic responses in the prefrontal cortex of mice were also demonstrated [20].

Studies using tDCS [21,22] reported changes in glutamate and GABA levels upon stimulation *via* MR spectroscopy (MRS), but information about the influence of tACS or TI on neurotransmitter balance is lacking. This information would be immensely relevant for a future therapeutic approach in motor disorders such as Parkinson’s disease, where disease progression is linked to changes of glutamatergic and GABAergic levels detectable by MRS [23].

Here, we report the first *in vivo* rodent study combining minimal-invasive TI with multiple MRI measurements. Following a systems approach, a finite-element-based digital twin of the rat head was established and used to optimize electrode placement and currents for targeting deep brain structures; the predicted stimulation effects were then examined *in vivo* on brain-wide neural patterns and connectivity using stimulation-evoked and resting-state fMRI, as well as on neurotransmitter balance with MRS.

## Results

### Computational Modeling and electrode optimization

A digital twin of the rat head, i.e., an anatomically realistic volume conductor model, was constructed from a T2-weighted anatomical MRI scan (see Methods and Supplementary Fig. 1). The scan was segmented into major tissue compartments, including scalp, skull, and brain. The brain compartment was further parcellated into anatomically defined regions ([24]; Fig. 1a). Based on this parcellation, the thalamus (Thal), in particular its ventrally located motor subdivision, was selected as the primary target region for subsequent stimulation.

**Figure 1:**
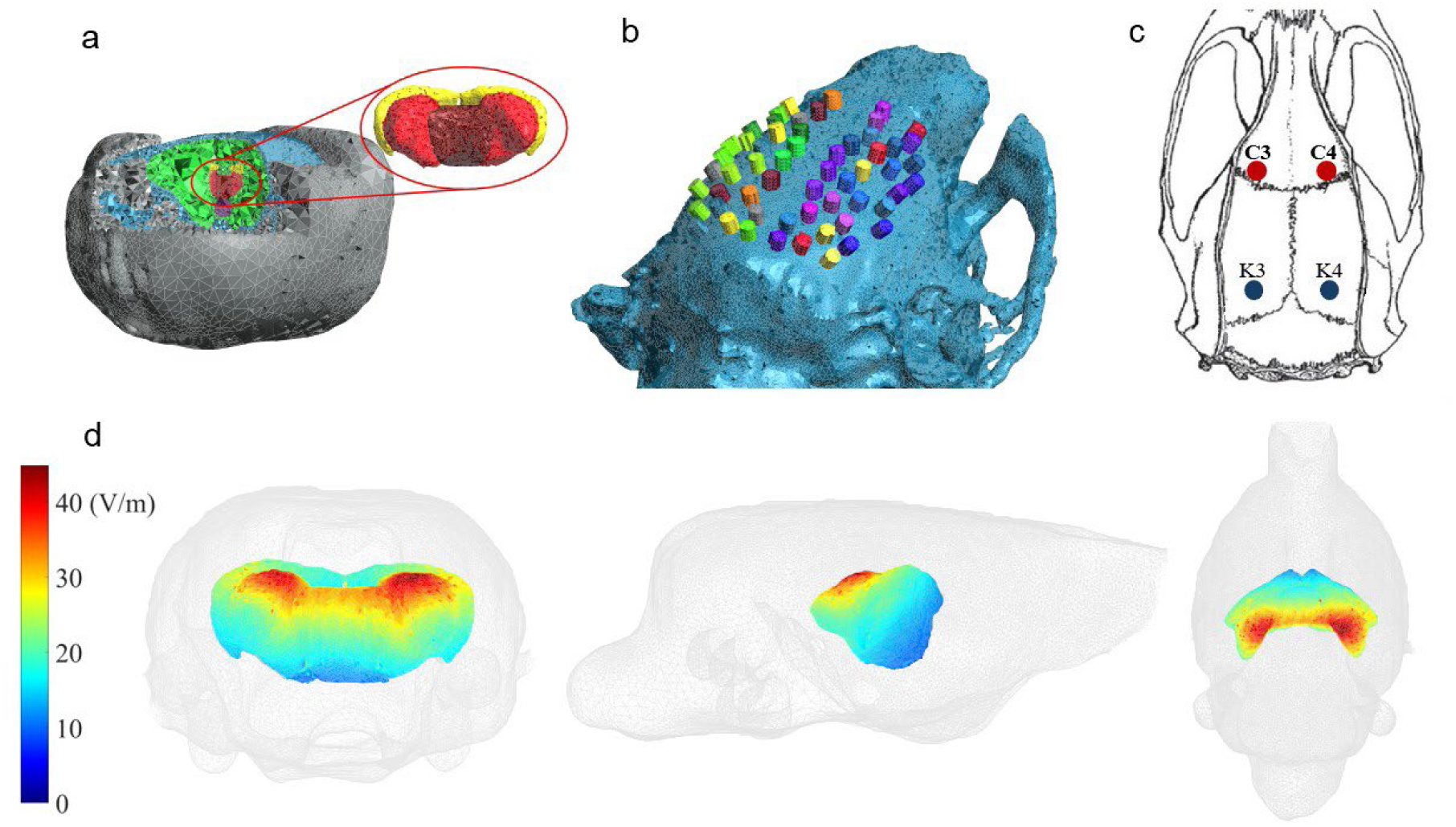
Model-based optimization of the electrode configuration for TI tACS. **(a)** Tetrahedral volumetric mesh of a rat head model showing tissue-specific domains used for electric field simulations in TI. Distinct regions include the scalp (gray), skull (light blue), hippocampus (yellow), thalamus (central thalamic region shown in dark red and adjacent thalamic subregions in red), hypothalamus (purple), and remaining brain tissue (green), with a magnified inset highlighting the primary stimulation target (thalamus) and an additional subcortical region examined in post-hoc analysis (hippocampus). **(b)** Placement of the 56 candidate electrodes on the skull of the rat head model. **(c)** Stereotactic coordinates of the rat‘s skull, adapted from the stereotaxic rat brain atlas [24] used for TI tACS. Red and blue circles indicate the positions of the first and second electrode pairs, respectively, used in the *in vivo* and simulation studies. **(d)** Modulation depth *E_MD_ of the optimized montage in deeper brain regions during TI stimulation, in V/m.* Notable modulations are observed in the predefined target regions including the thalamus. Color intensity represents field strength.

Several candidate electrode positions (Fig. 1b) were combined and evaluated to identify the optimal electrode pair configuration for stimulating the predefined target region. The optimization procedure identified the C3/C4 and K3/K4 electrode pairs as optimal (Fig. 1c) with an injection current amplitude of 1 mA applied to each pair. This configuration meets practical experimental constraints and is suitable for implementation in an *in vivo* TI experiment.

Simulation results demonstrated that TI achieves high modulation intensity within the Thal (Fig. 1d). Notably, electric-field modulation was also detected in the region between the electrode pairs, as well as in the underlying dorsal hippocampus (HIP); however, the modulation strength in the Thal was higher than that in the dorsal HIP. The target region thus showed the highest modulation depth among the deep ROIs, consistent with the optimization objective.

Although modulation strength decreased with increasing depth in the target area, the hypothalamus (HY) still exhibited a clearly detectable modulation, indicating that TI can effectively reach even deep subcortical structures.

Quantitative analysis of the simulated modulation depth further supports the spatial patterns observed in the electric field maps (Suppl. Tab. 2). Interestingly, Thal, the primary target region of interest, exhibited higher modulation depth than HIP; however, both regions displayed substantial modulation depth, indicating effective penetration of the electric field into deep brain structures. In the rat, TI thus reaches deep structures without a pronounced depth-dependent fall-off; regions close to the injection sites are, as expected from the model and consistent with the strong dependence of field strength on head size [14], not spared.

### Experimental setup

The experimental setup was largely adopted from Grossman and colleagues [11] and is illustrated in Figure 2, for all details see Supplementary Information.

**Figure 2:**
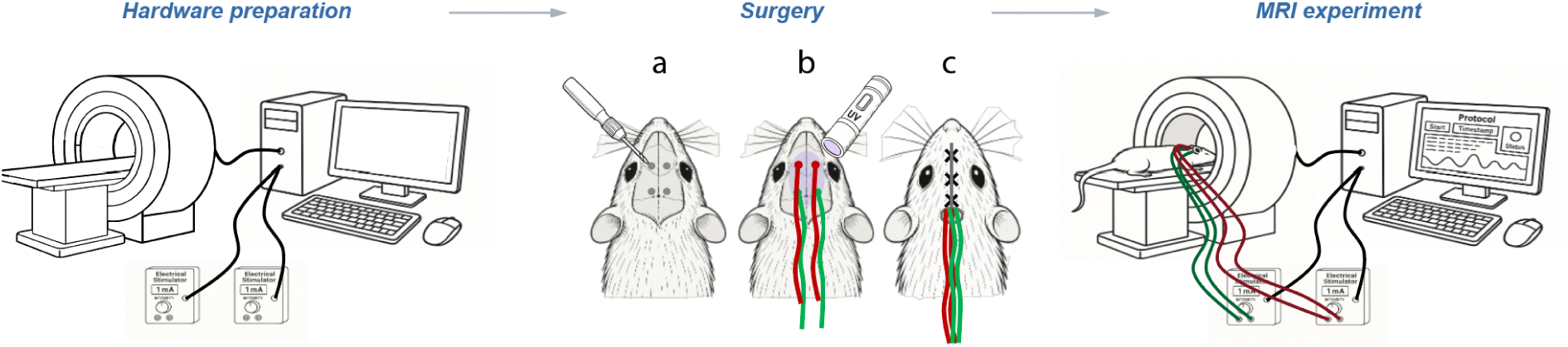
Overview of the consecutive procedure on the day of experiments as a timeline. Before animals underwent surgery, the TI setup was prepared by connecting the stimulators to a PC and the PC itself to the MRI scanner to synchronize both *via* a TTL pulse and tested for optimal performance (left). Major steps of the surgery on the anaesthetized rats include (a) cleaning and flattening of the exposed skull surface with a small dental drill (b) placing the two electrode pairs at positions C3/C4 and K3/K4, attaching them with glue and UV light, (c) followed by closing the surgical opening (middle). Afterwards, rats were immediately placed on an MRI-compatible cradle and the electrode cables were connected to the stimulators (right).

### BOLD fMRI

BOLD (blood oxygenation level dependent) activation pattern changes (Fig. 3) were recorded before and after TI at electrode locations C3/C4, K3/K4, with 1 mA current amplitude and f1=2000 Hz and f2=2010Hz as stimulation frequencies. The minimal-invasive stimulation induced bilateral BOLD changes in several brain areas, most importantly including predefined and simulated deep brain regions like the Thal (in particular ventrolateral, laterodorsal, posterior, and lateral posterior nuclei of the dorsal thalamus). As expected from the simulations, BOLD changes were also detected in the dorsal HIP as well as in regions close to the electrodes, such as anterior cingulate (ACA) and retrosplenial (RSP) cortex. In addition, cortical areas distant from the electrodes, particularly the infralimbic area (ILA) and visual cortex (VIS) as well as the amygdala (AMY) showed a BOLD signal change. Also, the basal forebrain and the subiculum (SUB) showed a BOLD signal change, but with lower activation than the other regions. The region-of-interest (ROI) analysis of one representative rat (Fig. 3b) showed a clear temporal profile in correspondence to the classical canonical hemodynamic response function (HRF) for cortical (ACA, VIS, RSP, ILA) and subcortical (HIP, AMY, Thal, SUB) areas, highlighting an initial dip at the beginning of the stimulation, followed by an increase up to 4 % (cortical) or 2 % (subcortical) BOLD signal change, and a recovery to baseline after eight to ten seconds in relation to the beginning of the stimulation. Interestingly, there is a visible difference in peak amplitude for cortical areas, located directly underneath the electrodes (RSP and ACA) and areas that are far more anterior (ILA) or posterior (VIS) which may reflect primary and secondary (via intracortical connections) stimulation effects. However, no signal gradient or lowered peak amplitude was observed for subcortical regions with respect to the depth of the regions, i.e., the dorsal HIP shows a comparable amplitude and timing as the more ventrally located Thal. This observation is exactly in line with the mean modulation depth of the computational simulations (see Suppl. Tab. 2). In addition, we detected a significant difference in the overall temporal profile (p < 0.05) when comparing mean curves of cortical and subcortical BOLD response (Fig. 3b, lower left panel). Individual time points, e.g. the maximum amplitude or the time-to-undershoot were not significantly different. However, the amplitude maximum was reached earlier in subcortical regions (4.5 ± 0.5 s) compared to cortical areas (5.5 ± 1.1 s). A bilateral negative signal change in the caudate putamen (CPu) did not exhibit a typical HRF-like temporal profile (Fig. 3b, lower right panel).

**Figure 3:**
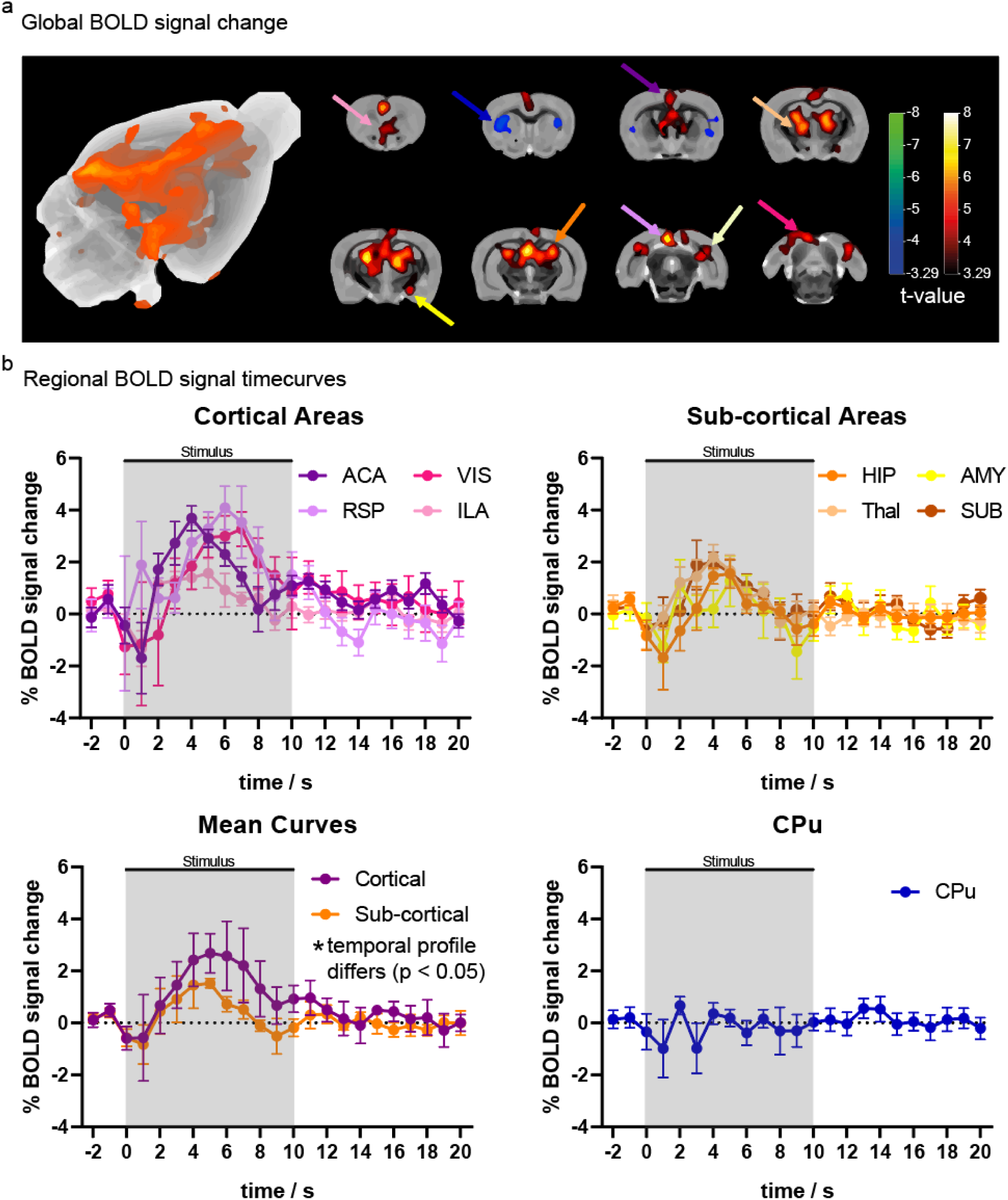
Group BOLD activation patterns of experimental rats during TI stimulation. **(a)** 3D glass brain view of the overall BOLD activation pattern (left panel) based on the group analysis of the experimental rats (n = 8) and individually selected slices of BOLD activation (right panel) from anterior to posterior overlaid with the anatomical rat template for a certain statistical threshold (p < 0.001, q < 0.05, t-value of ± 3.29). The color-coded arrows indicate the location of the ROI position for extraction of the signal time courses. **(b)** Exemplary temporal profiles in mean value ± standard deviation of selected regions-of-interest such as cortical regions (upper left), subcortical regions (upper right), the mean curve of cortical and subcortical regions (lower left) and the caudate putamen showing a negative signal change (lower right). A two-tailed unpaired student’s t-test (p < 0.05) was used to compute statistically significant differences of the mean curves for the overall temporal profile during the stimulation period.

In addition to the BOLD fMRI experiments presented above, we performed several control experiments. First, a series with a very low and thus most likely ineffective amplitude of applied current (0.1 mA) on rats (SHAM-group) demonstrated no positive BOLD signal changes at all (Suppl. Fig. S2). Second, to check if the applied electrical field itself induces magnetic susceptibility changes, a piece of lemon was scanned as a non-living proxy with the same settings. We found no signal change due to TI stimulation, even with a very loose p-value < 0.05 and no FDR-correction (Suppl. Fig. S3). Third, control measurements with a pickle were performed to investigate the effect of the applied electrical field on the alternating magnetic field (B1). Here, B1 maps (Suppl. Fig. S4) showed only small variances in amplitude at the edges of the pickle and an equally distributed variance for the phase of B1.

### resting-state fMRI

To investigate potential long-lasting effects of TI on functional connectivity, resting-state fMRI was performed before and after stimulation (Fig. 4). A significant decrease in overall correlation was observed in the second resting-state scan following the stimulation period, most prominently in the SHAM group (Fig. 4a). No significant global differences between groups were detected post stimulation. The distribution of correlation values (Fig. 4b) showed a general shift towards lower correlations in the SHAM group, consistent with the known decline in functional connectivity during prolonged anesthesia [25,26]. This reduction was more pronounced in the SHAM group (approximately +61.6% shift towards lower correlation values), whereas the experimental group exhibited a markedly smaller change (approximately −12.1%), suggesting that TI stimulation may mitigate this effect. Region-specific analysis (Suppl. Tab. 3) revealed higher correlations in SHAM animals prior to stimulation in the insular cortex (p < 0.01) and cerebellar regions (p < 0.0001). The fimbria of the hippocampus showed higher correlations in SHAM both before (p < 0.005) and after stimulation (p < 0.01), while the entorhinal cortex differed only post stimulation (p < 0.005).

**Figure 4:**
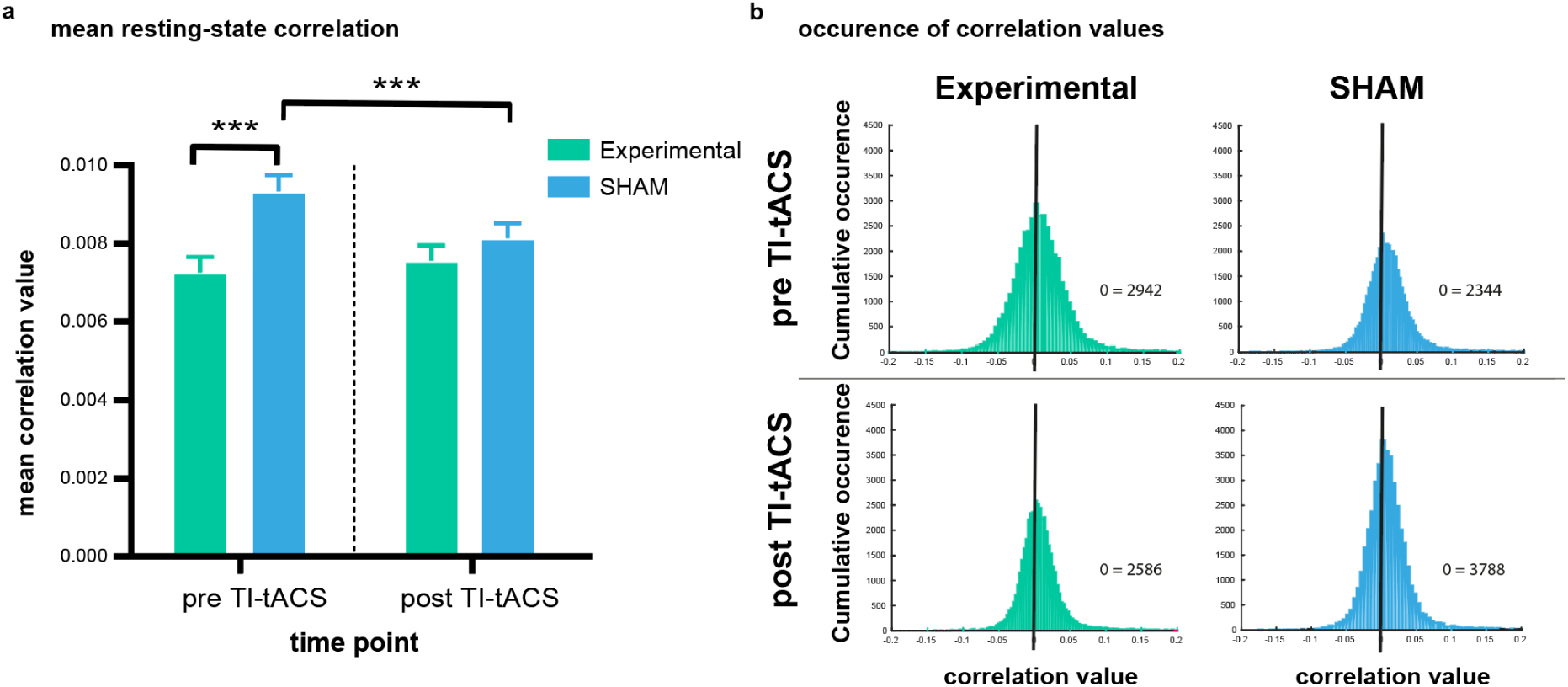
Resting-state correlation at different time points. **(a)** Correlation values averaged across 234 brain regions of the Sigma rat brain atlas [27] for the two different time points of all experimental rats with TI stimulation of 1 mA (n = 8) and SHAM-treated rats (0.1 mA, n = 5). A repeated-measures ANOVA with Bonferroni correction (p < 0.05) was performed to test for statistical significance. **(b)** Histograms of the above calculated Pearson correlation coefficients for both time points and groups showing the occurrence of every correlation value. Correlation values of “0” were summed up. The black vertical line is located at a correlation value of zero.

### MR spectroscopy

The MR spectroscopy (MRS) revealed further insights into metabolic changes of neurotransmitter levels. The spectroscopy voxel was placed within the Thal, the simulated target region (Fig. 5a). A representative spectrum of a rat pre, during and post stimulation is illustrated in Figure 5b. Here, metabolic changes for the total amount of Glx (glutamine + glutamate) are already identifiable in the raw data. The total amount of Glx and GABA (y-aminobutyric acid) was extracted for all rats and tested for statistical significance. No significant difference in neurotransmitter levels was found when comparing the different time points (Fig. 5c). However, trends between pre-stimulation and post-stimulation were identified. The experimental group showed an increase in Glx during TI and no change in GABA. This leads to an increase for the ratio Gx/GABA during stimulation. No changes were observed in other metabolites, such as NAA (N-acetyl-aspartate), supporting the reliability of the measurements.

**Figure 5:**
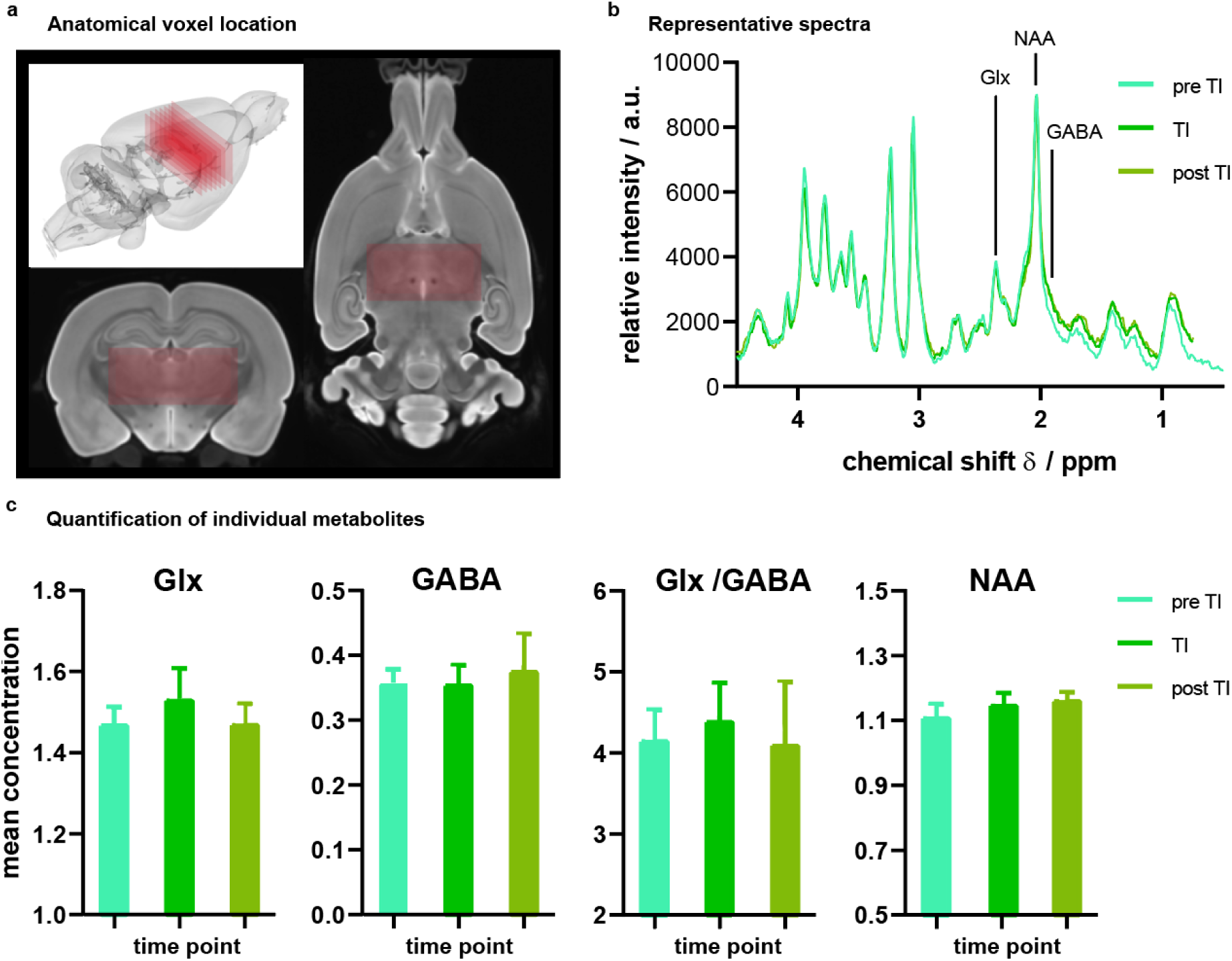
MR spectroscopic results for all three time points. **(a)** Visualization of the voxel position as a red box within a 3D glass brain, a 2D frontal and a 2D horizontal view. **(b)** An exemplary MR spectral overlay of one experimental rat pre, during and post stimulation. The black vertical lines mark the chemical shift of the metabolites of interest (Glx, GABA and NAA). **(c)** Quantification of the metabolic changes in mean values ± s.e.m. for all three time points: pre/ during/ post stimulation (number of rats = 4/4/3) for Glx (left), GABA (2nd left), Glx/GABA ratio (3rd left) and NAA (right). Differences between the three experimental timepoints (pre, during, and post stimulation) were assessed using a linear mixed-effects model with the subject as a random effect to account for repeated measurements (p < 0.05).

## Discussion

Our results confirm the feasibility of applying transcranial temporal interference stimulation (TI) *in vivo* in rats, establishing the method as a viable approach for future experimental and translational research. The electrode configuration was optimized on a computational model of the rat head to target deep structures, in particular the thalamus, and the experimental setup and surgical procedure were adapted to apply TI during fMRI and MRS. Here, we clearly identified cortical and subcortical regions with altered BOLD signal patterns and functional connectivities (FC) upon stimulation. A strong BOLD change was detected in the predicted targets, in line with the modulation depth predicted by the digital twin (Suppl. Tab. 2); the fMRI readout thus closes the loop between model-based design and experimental validation. FC varied in brain regions with a BOLD response, but also in regions which are functionally connected with these BOLD activated regions. In addition, the thalamic neurotransmitter balance indicates an increase towards excitatory glutamate during TI for rats showing BOLD response.

The basic idea of applying TI as a therapeutic approach is to target well-defined deep brain regions, without stimulatory effects on the overlying cortex. In our study, we have detected BOLD responses in predefined deep brain target regions from our simulations, such as the thalamus, the hippocampus, and the amygdala, as well as in cortical areas not underlying the electrodes. Unlike in the mouse study [11], cortical activation under the electrodes was both predicted by the model and observed by fMRI, which we attribute to the small rat head and the skull-mounted electrodes rather than to a failure of TI focality.

We detected changes in FC with rs-fMRI after stimulation, especially in the insular cortex. Interestingly, this structure did not show a BOLD response to the stimulation, but it is functionally connected to the amygdala [28], a region with increased BOLD. This connectivity difference is observed when comparing the resting-state signal before stimulation, but not afterwards. A significant difference in FC was also observed for the fimbria of the hippocampus, being a part of the hippocampal area, again a region with altered BOLD signal change and one of the target regions. Together with the differences in peak amplitudes of BOLD activations in cortical areas located directly beneath the stimulation electrodes (RSP and ACA) and in more distant regions (ILA and VIS), we interpret the observed changes in FC as reflecting primary versus secondary effects of stimulation mediated through the respective neuronal connections.

Finally, MR spectroscopic results shed light on the neurotransmitter balance alterations due to TI. Although the ratio of Glx/GABA was not significantly different between the groups, a clear trend towards excitatory glutamate was observed. So far, the spectroscopic voxel was large in dimensions and placed centrally in the thalamus, a structure with stimulus-induced BOLD response, but the thalamus did not respond globally in the same way to the stimulation. This leads to higher variability and, therefore, to less distinguished neurotransmitter quantification. Chemical shift imaging (CSI) would allow a simultaneous spectroscopy of different brain regions and the identification of the most prominently affected thalamic nuclei.

The success of TI may be strongly related to the individual brain state, as it is discussed elsewhere for tACS in humans [29,30]. Although we standardized the experimental procedure, individual stress levels or the anesthetic depth may have been the reason why some animals did not show stimulus-induced BOLD changes at all. Likewise, anesthesia affects neurotransmitter levels, and medetomidine, as an A2-agonist, modulates glutamatergic and GABAergic release [31], which impacts the MR spectroscopy results. An awake-MRI approach in well-habituated animals with permanently attached electrodes would make anesthetic effects negligible [32].

On the modeling side, the digital twin was derived from a single representative animal, with isotropic literature conductivities and without a separate CSF compartment. Individualized computational models that are updated with fMRI and MRS readouts are the natural next step towards personalized and, eventually, closed-loop TI [15,33].

In conclusion, our findings demonstrate the feasibility of TI for *in vivo* use in rats, establishing a methodological basis for future studies investigating its translational potential in neurological and neuropsychiatric research. Future work should aim to refine surgical and experimental parameters and translate these findings towards clinical applications or disease models to bridge our and other animal research with human studies.

## Materials & Methods

### Computational modeling and simulation

#### Volume Conductor Modeling

Raw DICOM images (Suppl. Fig. S6) from a high-resolution anatomical MRI dataset were converted to NIfTI format and preprocessed through bias correction, registration, and segmentation. Intensity inhomogeneities were addressed using N4 bias field correction, followed by manual alignment. Brain, skull, and scalp tissues were segmented semi-automatically in 3D-Slicer v5.10 [34]. To replicate the experimental setup, the upper scalp was removed, and the skull surface thinned. The hippocampus, thalamus, and hypothalamus were manually delineated as separate regions of interest (ROIs), yielding six regions comprising tissue classes and anatomical ROIs. Segmentations were refined and merged into a multi-label volume. Tetrahedral meshes with tissue-specific conductivities were generated in SimNIBS v4.6 [35], which served as the basis for subsequent electric-field simulations.

#### Forward Modeling and Optimization of TI tACS

The induced electric potential within the conductive medium was computed by solving the generalized Laplace equation, and the electric field was obtained as its spatial gradient. The resulting field distributions from the two sources were combined to estimate the modulation depth [35,36]. The heterogeneous rat-head geometry was solved numerically using the Finite Element Method (FEM) in SimNIBS v4.6 [35,36]. A leadfield-based forward model mapped electrode currents linearly to intracranial electric fields. Optimization was performed to identify two bipolar electrode pairs and corresponding current amplitudes that maximize the modulation depth ratio within a target region relative to off-target areas, subject to current conservation and safety constraints, following the leadfield-based optimization approach of our earlier work [13,37]. Further implementation details are provided in the Supplementary Material.

### Experiments

#### Experimental setup

The TI setup (for technical details of all used hardware devices, see Supplementary Table 4) adapted from Grossman et al. 2017 [11] comprised two LabView-controlled stimulators delivering 1 mA (SHAM: 0.1 mA) through two pairs of Ag/AgCl electrodes. Their frequencies of 2.00 kHz and 2.01 kHz produced a 10 Hz interference difference.

#### Animal handling and surgery

All rats used for this study were housed in a 12/12-hour light/dark cycle with food and water *ad libitum*. All experiments were performed according to the Directive of the European Communities Parliament and Council on the protection of animals used for scientific purposes (2010/63/EU) and were approved by the animal care committee of Sachsen-Anhalt, Germany (42502-2-1653 LIN). Male Wistar Han wild-type rats (16 weeks, 388 ± 35 g) were used. Animals were anaesthetized with isoflurane (3 vol% in normal breathing air for anesthesia induction, 1.5 vol% for maintenance). Rats were placed on a heating pad for body temperature stabilization at 37 °C. Under microscopic control, the scalp was opened and the skull surface cleaned and flattened with a drilling machine to maximize bone contact of the electrodes. Ag/AgCl electrodes were attached using UV glue at the modelled positions (C3/C4, K3/K4). The incision was sutured, leaving a small hole for cable exit.

For MRI measurements a tail vein catheter filled with a 1:10 medetomidine/NaCl solution was placed. Afterwards, animals were transferred to the MRI scanner and placed on a rat cradle with a bite bar and nose tube, allowing gentle head fixation and inhalation gas delivery. Vital parameters of the rats (breathing rate 80 ± 10 bpm, body temperature 37 °C) were observed via a MR-compatible monitoring system. An air heater system delivered warm air to keep the body temperature stable.

#### MRI experiments of the rats

MRI data were acquired using a 9.4 T Bruker AVANCE III small-animal MRI scanner (20 cm bore) equipped with a BGA12S gradient system. Data acquisition and reconstruction were controlled by ParaVision 7 software. All parameters for the following sequences are listed in Supplementary Table 5. The simulation dataset was acquired from a 20-week-old male rat, anaesthetized as above, using a 075/40 ^1^H transmit/ receive volume coil. A 2D T2-weighted TurboRARE anatomical reference sequence was acquired was followed by a high-resolution 2D T2 TurboRARE (Suppl. Fig. S1).

For all functional and spectroscopic measurements, a 112/086 ^1^H volume resonator was used for transmitting and a ^1^H single-loop surface coil (ID20) for receiving signal. Initial anesthesia was switched to medetomidine (i.v., 0.05 mg/kg body weight/hr). Then, a ^1^H Localized Shim placed in the thalamus (for exact location see Fig. 4) was used to minimize inhomogeneities and to perform MR spectroscopy. A ^1^H-PRESS sequence was acquired. The fMRI experiments started after the pre-stimulation MRS, 60 minutes after the start of medetomidine administration, and were performed with a 2D GE-EPI sequence. One resting-state scan and a stimulation-evoked scan were acquired. The stimulation paradigm was presented in a block design (120 s baseline, 20 repetitions of 10 s stimulation, and 20 s rest). Afterwards, a second resting-state scan was acquired, followed by two MRS scans, one with a stimulation (20 repetitions, 10s stimulation, 10s rest) and one without. All MRI scans were exported to DICOM format *via* the ParaVision 7 software, spectroscopies to ASCII format via Topspin 3.2.

#### MRI experiments of the phantoms

Two phantoms were scanned with the rat hardware setup and Ag/AgCl electrodes taped to their surface. A lemon was used to assess stimulation-induced susceptibility artefacts with the *in vivo* evoked fMRI paradigm. A pickle was used to assess B1 inhomogeneities using two maps without and one with stimulation. Datasets were exported to DICOM.

#### MR analysis

All stimulation-evoked fMRI datasets (n = 16 rats in the experimental group and n = 7 animals in the SHAM group) were analyzed with BrainVoyager 20.6 software, starting with preprocessing, including removal of the first 10 volumes, slice-time correction, 3D motion correction, high-pass filtering, and a 3D spatial smoothing of two pixels. Afterwards, individual scans were registered to our anatomical MR image of the SIGMA rat brain atlas [27]. A general linear model (GLM) approach was used to investigate BOLD pattern changes with an HRF (time-to-response-peak = 2 s, time-to-undershoot-peak = 10 s). Eight out of 16 experimental rats showed no BOLD signal change and were thus excluded. Technical failure was considered unlikely because MRI acquisition was error-free, stimulator output and electrode continuity were verified before and after scanning, and post-scan inspection excluded adhesive-related contact insulation. Anesthesia-related response suppression remained a possible explanation. For the group analysis, the statistical threshold was set to a p-value < 0.001, with FDR-correction (q < 0.05). With an event-related averaging, mean signal time courses ± standard deviation for regions showing BOLD signal changes were extracted. Group analysis images were exported to NIfTI format, and ROI time courses to text format. Images were imported into MRIcroGL 4.6 and overlaid with the rat brain template.

All resting-state datasets (n = 8 in the experimental group; n = 5 in the SHAM group) were analyzed with AIDAmri 2.0 (https://github.com/Aswendt-Lab/AIDAmri) [38] and AIDAconnect 1.1 (https://github.com/Aswendt-Lab/AIDAconnect) [39] using adapted FSL 6.0 algorithms [40–42] for bias correction [43], brain extraction [44], 3D motion correction [45,46], and slice-time correction. All scripts were revised to fit the Sigma Rat Brain Atlas [27]. Group correlation matrices and their mean values based on the z-transform of the Pearson coefficient were calculated and used to compare the two experimental time points and groups globally (repeated-measures ANOVA, p < 0.05, Bonferroni-corrected) and region-specific (multiple t-tests, p < 0.05, Holm-Sidak-corrected). MR spectroscopy data were analyzed with LCModel 6.3 [47,48] and the basis set for PRESS sequences (echo time = 15 ms at 9.4 T, spectral width from 4 to 1 ppm) modeling the measured spectrum as a linear combination of metabolite basis spectra with a spline baseline. Concentrations were estimated from fitted scaling coefficients, water-scaled, and normalized to total creatine. Mean values ± standard error of the mean were calculated for Glx, GABA, NAA, and Glx/GABA. Differences between the three experimental timepoints (pre, during, and post stimulation) were assessed using a linear mixed-effects model with the subject as a random effect to account for repeated measurements (p < 0.05). The B1 maps from the pickles were analyzed with a custom-written MATLAB script, building the differential amplitude and phase images for the different experimental conditions. The stimulation-evoked fMRI data from the lemon was analyzed in the same manner as the *in vivo* data (p < 0.05, no FDR-correction). All statistical tests and plots were performed with GraphPad Prism 8.0.1.

## Acknowledgements

This study was supported by the federal state of Saxony-Anhalt and the European Regional Development Fund (ERDF) in the Center for Behavioral Brain Sciences (CBBS, ZS/2016/04/78113). MA and AK acknowledge financial support by the Deutsche Forschungsgemeinschaft (DFG, German Research Foundation): project ID 431549029–SFB 1451 project C01 and the Chan-Zuckerberg-Initiative funded project “COMULISglobe” (reference number 2023-321161).

We thank Dr. Michael Lippert for technical support in LabView programming, as well as Janet Stallman and Anja Gürke for excellent technical assistance.

## Author Contributions

RF, PR, and EB acquired funding for the project and conceived of the experiments, with additions to experimental design from PW, KA, and FA. Hardware devices for the experimental setup were established by FA. The computational model was designed by KA and RF. Pilot tests were performed by PW, FA and AM. *In vivo* experiments were performed by PW and EB, phantom experiments by PW. Data was analyzed by PW, KA, and DM in consultation with AK, RF, EB, TZ and MA. Results and interpretation were done by PW, KA, EB, and RF, and discussed with all authors. PW and KA wrote the manuscript with input from all authors.

## Supplementary Information

### Simulation of transcranial temporal interference stimulation

TI tACS is a noninvasive technique that enables deep and spatially selective brain stimulation by superimposing two high-frequency sinusoidal currents with slightly different frequencies (*f*_1_≈*f*_2_) [11,34]. Although each current alone oscillates at frequencies generally considered too high to effectively trigger neuronal firing, their interaction within brain tissue may generate a low-frequency amplitude modulation that could influence neural activity in targeted regions [11,27]. The induced electric potential *φ*(*x*) in the conductive domain is governed by the generalized Laplace equation, −*∇* ⋅ (*σ*(*x*)*∇φ*(*x*)) = 0, where *σ*(*x*) denotes tissue conductivity [35,36]]; the corresponding electric field is given by 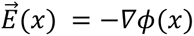.

The superposition of the resulting fields 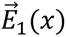 and 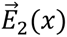 produces modulation at the beat frequency |*f*_1_ *f*_2_|, and neural activation is quantified by the modulation depth across projection directions n̂,

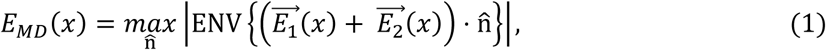

evaluated using the closed-form maximum-envelope expression [11,12], as implemented in SimNIBS. When 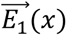 and 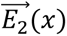 are collinear, this expression reduces to

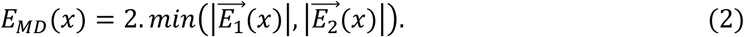

Due to the complex geometry of the rat head and heterogeneous, anisotropic tissue properties, analytical solutions for the electric potential and resulting modulation depth are not feasible, necessitating numerical approaches based on realistic volume conductor models.

### Forward model approximation

To solve the Laplace equation numerically and compute the electric field, the conductive domain was discretized into tetrahedral elements using the Finite Element Method (FEM). Tissue-specific conductivities are assigned to each compartment, and the electric potential *φ*(*x*) and field 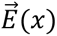 are computed in SimNIBS v4.6 [35,36]. However, repeatedly solving the full FEM problem for different electrode configurations is computationally expensive, especially in optimization tasks. To address this, a forward-model approximation based on a leadfield matrix *LεR*^3*n*×*m*^ was used, where *m* is the number of electrodes and *n* the number of mesh nodes; this matrix linearly maps electrode currents 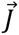 to intracranial electric field components via 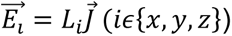. This relationship, derived from the superposition principle, enables fast evaluation of the electric field and efficient computation of the modulation depth at each node, providing a scalable framework for TI tACS optimization.

### Leadfield-based optimization of transcranial temporal interference stimulation

The optimization problem seeks to identify two bipolar electrode pairs, each operating at a distinct carrier frequency—from *m* candidate electrodes, along with their injected current amplitudes, to maximize the stimulation effect. Electrode selection is encoded by binary vectors 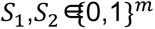, while the corresponding currents are represented by continuous vectors *I*_1_,*I*_2_⊠*R^m^*. Current amplitude was biased toward the safety-permitted maximum through the repair operator, which reset sub-maximal current pairs to *I_max_* unless they improved the focality ratio. The objective is to maximize the ratio between the average modulation depth in the target region(*Ω^t^*) and that in the off-target region (*Ω^ot^*):

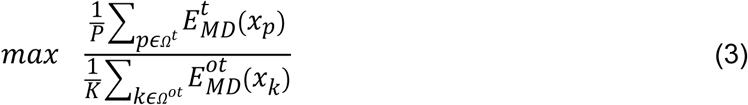

where, *P* and *K* denote the number of mesh nodes in the respective regions. The optimization is subject to current conservation 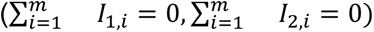, safety bounds(|*I*_1,*i*_|, |*I*_2,*i*_| ≤ *I_max_*) and bipolar constraints requiring exactly two active electrodes per frequency 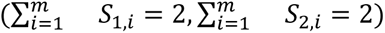. Due to the combination of discrete electrode selection and continuous current optimization, the problem is a mixed discrete–continuous optimization task. To efficiently handle this hybrid structure, a genetic algorithm is employed that jointly optimizes electrode configurations and current amplitudes for focal TI tACS targeting [13,37].

### Supplementary Figures

**Supplementary Figure S1:**
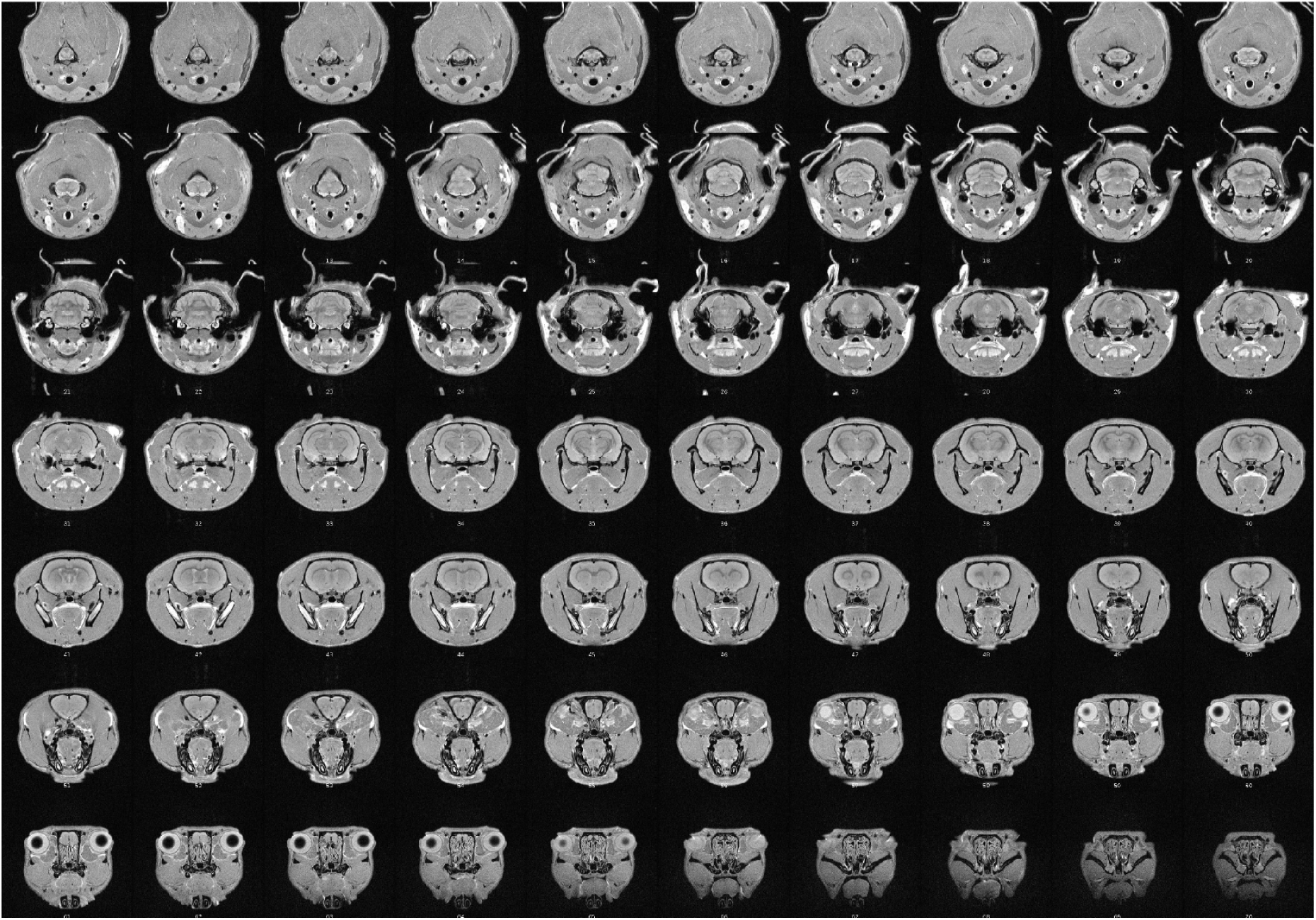
High-resolution anatomical MR images of the rat head serving for the computational simulations. Slices of the T2 TurboRARE scan are arranged from posterior (upper left) to anterior (lower right) direction.

**Supplementary Figure S2:**
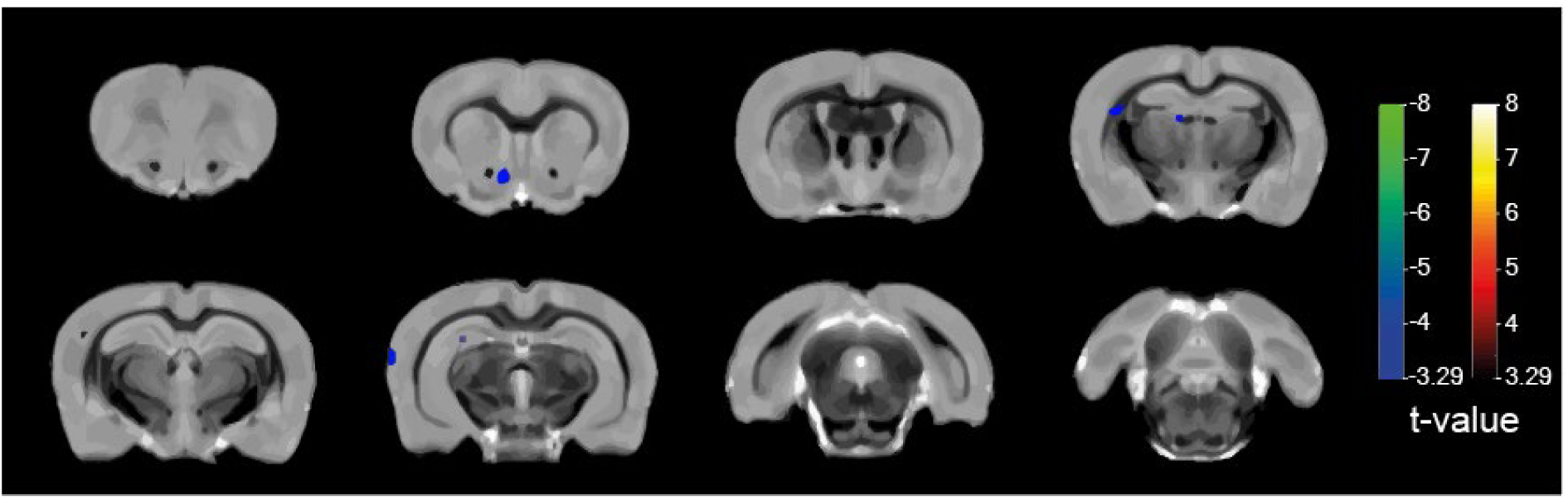
Group BOLD activation patterns of rats treated with a very low current (0.1 mA, SHAM group). Individually selected slices showing BOLD pattern changes from anterior to posterior overlaid with the anatomical rat brain template for animals (n = 7), which underwent a TI stimulation with a reduced and thus ineffective current (0.1 mA). Note the already less strict statistical threshold (p < 0.001, no correction for multiple comparison).

**Supplementary Figure S3:**
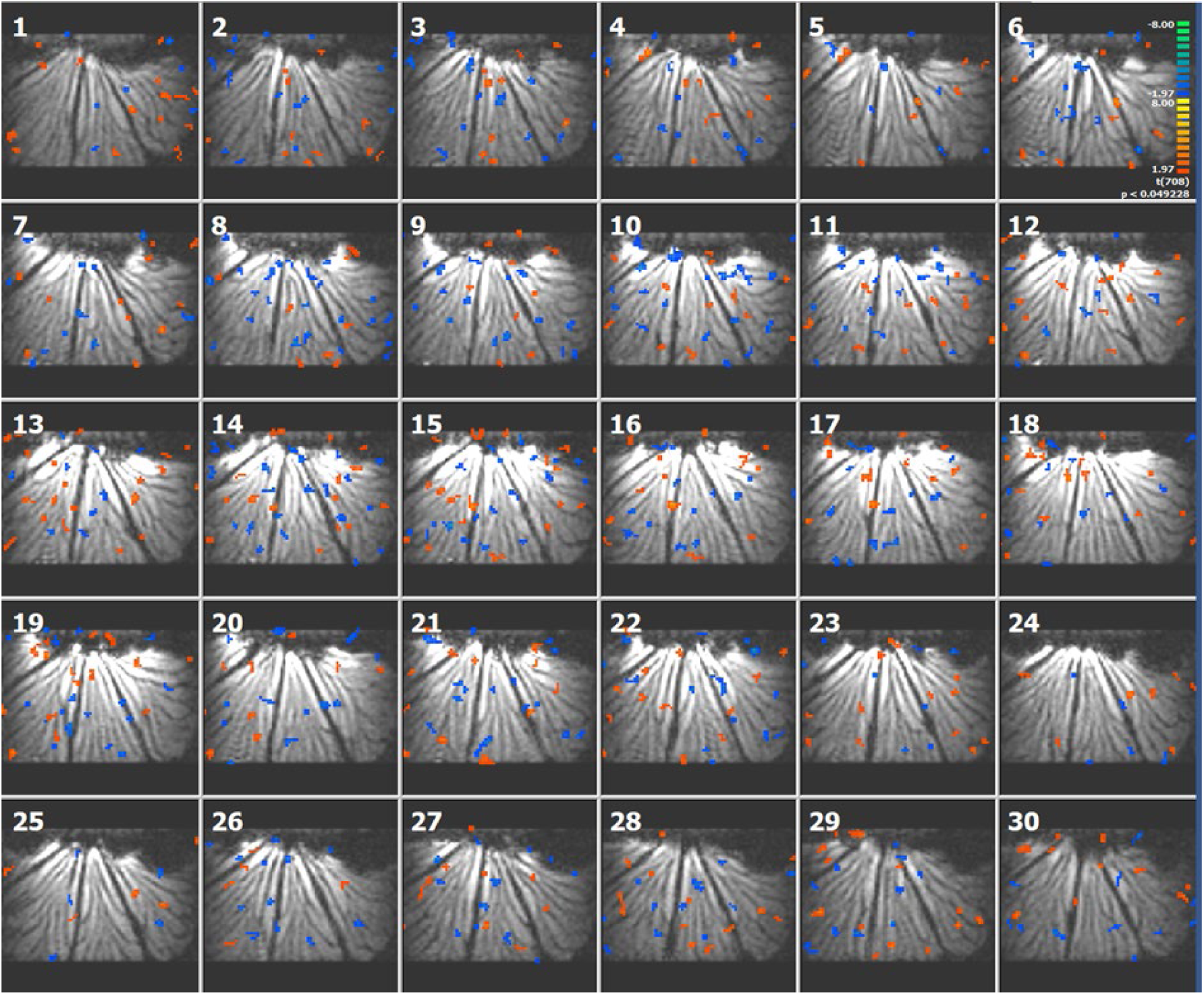
TI approach applied to a lemon. Voxel map of a single half of a lemon stimulated with TI (1 mA) demonstrated for 30 individual slices distributed all over the piece of fruit. The statistical threshold was set to p < 0.05, without correction for multiple comparisons.

**Supplementary Figure S4:**
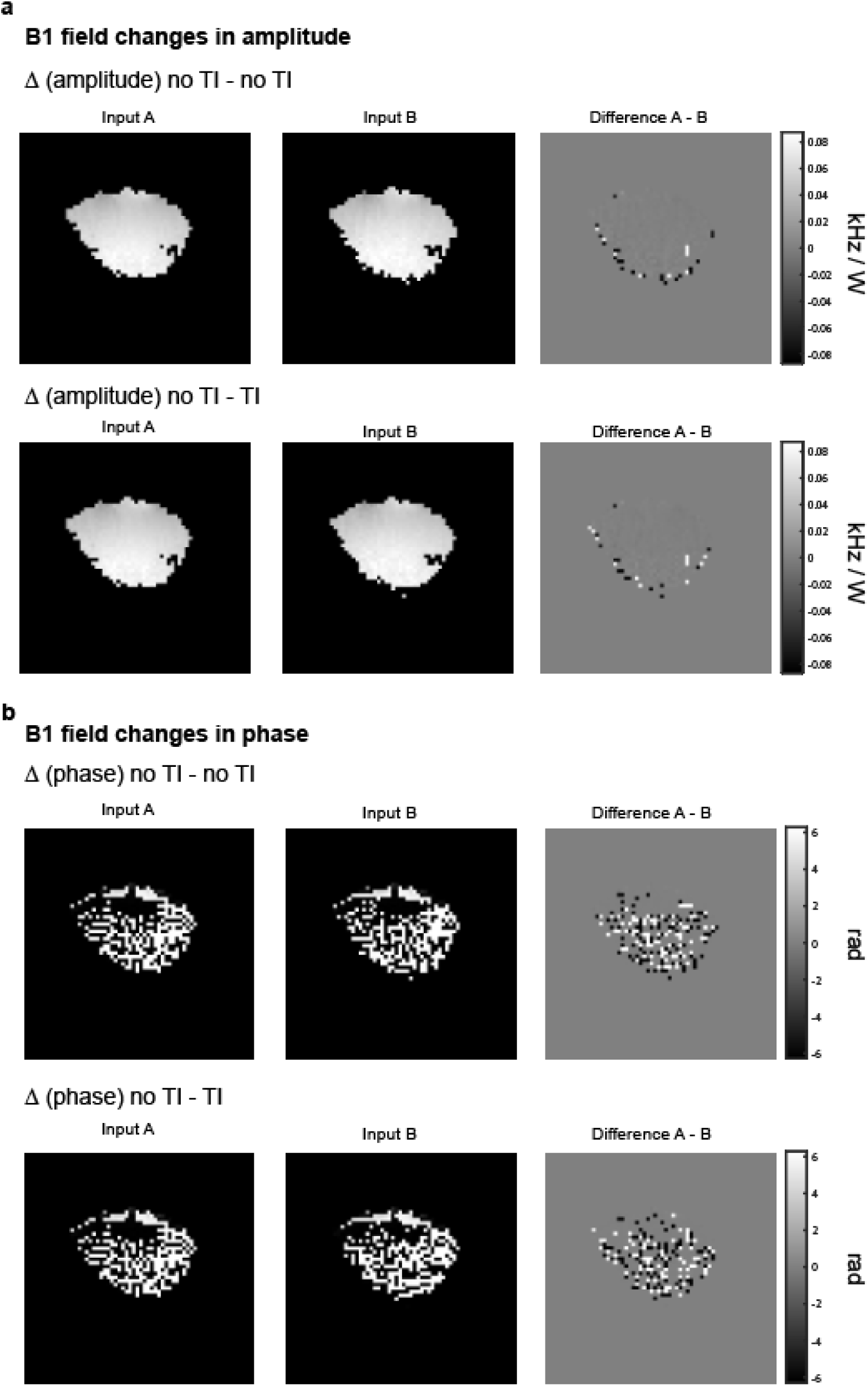
Differential B1 mapping of a pickle as control experiment. B1 amplitude maps in kHz/W **(a)** and B1 phase maps in rad **(b)** under different conditions (with TI and without TI). The original images are displayed in the left and middle row, the corresponding difference image on the right. The general hardware variability is detected when comparing two B1 maps without an additional electrical field and the variance induced by TI when comparing a B1 map without stimulation and one with stimulation.

**Supplementary Figure S5:**
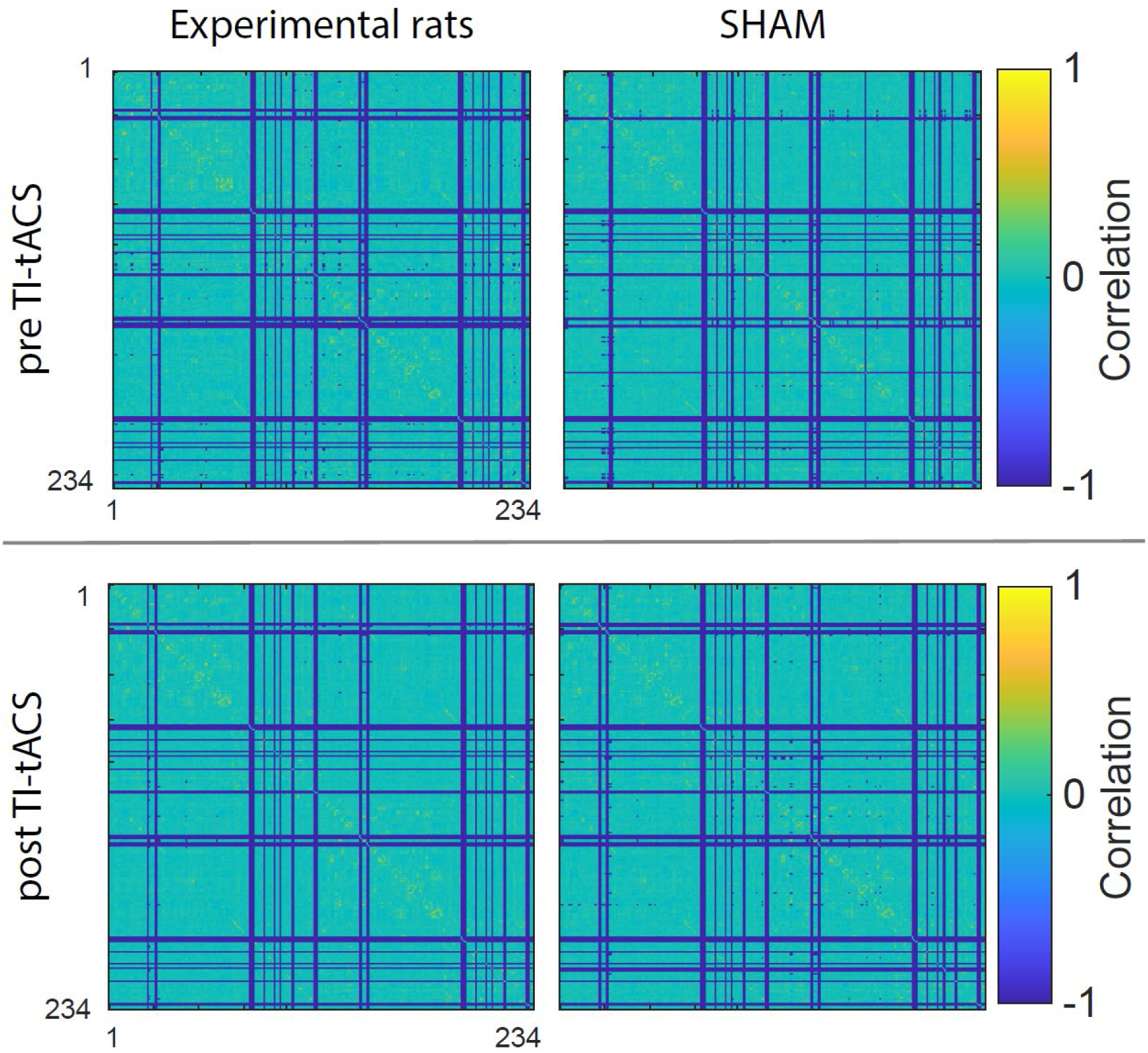
Resting-state correlations before and after TI stimulation in experimental and control rats. Correlation matrices of all 234 brain regions (structures and their numbering can be found the literature [27]) of the Sigma rat brain atlas [27] for the two different groups (experimental rats n = 8, SHAM rats n = 5) pre TI (upper panel) and post TI (lower panel). The z-transformation of the Pearson correlation coefficient is visualized ranging from -1 to 1, where 1 stand for a perfect positive correlation and -1 indicates a perfect anti-correlation. Blue vertical or horizontal lines indicate a region, where no correlation could be computed.

### Supplementary Tables

**Supplementary Table 1:** Electrical conductivities of tissue compartments in the rat head model. Values are presented in Siemens per meter (S/m).

| Tissue Name | Conductivity Value (S/m) |
| --- | --- |
| Scalp | 0.465 |
| Skull | 0.01 |
| Hippocampus | 0.276 |
| Thalamus | 0.276 |
| Hypothalamus | 0.276 |
| Remaining part of the brain | 0.2 |

**Supplementary Table 2:** Mean and maximum modulation depth *E*_MD_ in regions of interest during TI stimulation.

| Region of Interest | Mean $E_{MD}$ (V/m) | Max $E_{MD}$ (V/m) |
| --- | --- | --- |
| Hippocampus | 15.19 | 44.05 |
| Thalamus | 17.66 | 46.84 |
| Hypothalamus | 7.45 | 20.71 |

**Supplementary Table 3:** Region-specific significance of correlation values derived from resting-state fMRI between the individual groups tested with a multiple t-test and corrected for multiple comparison using the Holm-Sidak method (p-value < 0.05). The group comparison is shown in the left column and the regions according to the Sigma rat brain atlas with a significant difference are shown in the middle column with the corresponding p-value (right column).

| Groups | Region | p-value |
| --- | --- | --- |
| pre TI tACS<br>Experimental vs SHAM | L_Agranular_Dysgranular_Insular_Cortex | <0.01 |
|  | R_Molecular_Layer_of_the_Cerebellum | <0.0001 |
|  | R_Posterior_Agranular_Insular_Cortex | <0.01 |
|  | R_Fimbria_of_the_Hippocampus | <0.005 |
| post TI tACS<br>Experimental vs SHAM | L_Fasciculus_Retroflexus | <0.001 |
|  | R_Entorhinal_Cortex | <0.005 |
|  | L_Fimbria_of_the_Hippocampus | <0.01 |

**Supplementary Table 4:** List of hardware and software suppliers including all necessary details.

| Hardware & Software | Supplier and material information |
| --- | --- |
| TI tACS stimulator | Analog Stimulus Isolator Model 2200, A-M Systems Inc., USA |
| LabView | Version 2016, National Instruments, USA |
| Electrodes | 4 mm diameter, 100 cm lead wired, 1.5 mm touch-proof connector, NeuroSpec AG, Switzerland |
| Dental drilling machine | MF-Perfecta, W&H, Austria |
| Animal provider | Charles River Laboratories International Inc., USA |
| UV glue | Admira Fusion Flow, VOCO GmbH, Germany |
| Catheter tubes | inner diameter of 0.23 – 0.33 mm, Instech Laboratories, USA, and a cannula with 0.3 x 13 mm size, Braun Sterican, B. Braun, Germany |
| Medetomidine | 1mg/mL, Orion Corporation, Finland |
| NaCl solution | 0.9%, B. Braun, Germany |
| MR-compatible monitoring system | Model 1030, Small Animal Instruments, Inc., NY, USA |
| MR-compatible air heater system | Small Animal Instruments, Inc., NY, USA |
| MRI system | Bruker Biospin GmbH & Co. KG, Germany |
| ParaVision 7 | Bruker Biospin GmbH & Co. KG, Germany |
| Topspin 3.2 | Bruker Biospin GmbH & Co. KG, Germany |
| BrainVoyager 20.6 | Brain Innovation B.V., The Netherlands |
| MRicroGL 4.6 | https://www.nitrc.org/projects/mricrogl |
| MATLAB | MATLAB 2023a MathWorks, USA |
| GraphPad Prism | Version 8.0.1, https://www.graphpad.com/ |

**Supplementary Table 5:** Settings for the parameters of the individual acquired MRI/MRS sequences.

| Sequence | Parameters |
| --- | --- |
| T2 TurboRARE<br>(for anatomical reference) | Echo time = 24 ms<br>Repetition time = 2200 ms<br>Matrix = 355 x 234<br>Number of slices = 25<br>Spatial resolution = 0.1 x 0.1 x 0.8 mm<br>Number of averages = 6<br>Total scan time = 5 minutes |
| T2 TurboRARE<br>(for anatomical high-resolution scan) | Echo time = 6 ms<br>Repetition time = 2400 ms<br>Matrix = 256 x 256<br>Number of slices = 64<br>Spatial resolution = 0.12 x 0.12 x 0.5 mm<br>Number of averages = 5<br>Total scan time = 11 minutes |
| <sup>1</sup>H PRESS<br>(for MR spectroscopy) | Echo Time = 16 ms<br>Repetition time = 2500 ms<br>Voxel size = 6 x 4 x 4 mm<sup>3</sup><br>Water suppression method = VAPOR<br>Number of reference scans = 1<br>Total scan time = 10 minutes |
| 2D GE-EPI<br>(for functional MRI) | Echo time = 15 ms<br>Repetition time = 1000 ms<br>Matrix = 75 x 50<br>Number of slices = 25<br>Spatial resolution = 0.4 x 0.4 x 0.8 mm<br>Number of dummy scans = 5<br>Number of repetitions = 720<br>Total scan time = 12 minutes |
| B1 map | Echo time = 2 ms<br>Repetition time = 6 ms<br>Number of slices = 1<br>Spatial resolution = 0.75 x 0.75 x 1.5 mm<br>Number of averages = 1<br>Total scan time = 8 s |

## Abbreviation List

ACA: Anterior Cingulate Area AMY - Amygdala
BOLD: Blood Oxygen Level Dependent
bpm: breaths per minute
CPu: Caudate Putamen
CSI: Chemical Shift Imaging
DBS: Deep Brain Stimulation
FC: functional connectivity
FEM: Finite Element Model
fMRI: functional Magnetic Resonance Imaging
GABA: y-aminobutyric acid
GE-EPI: Gradient Echo – Echo Planar Imaging
GLM: General Linear Model
Glx: glutamine + glutamate
HIP: Hippocampus
HRF: Hemodynamic Response Function
HY: Hypothalamus
ILA: Infralimbic area
MR: Magnetic Resonance
MRI: Magnetic Resonance Imaging
MRS: Magnetic Resonance Spectroscopy
NAA: N-acetylaspartate
Ppm: parts per million
PRESS: Point RESolved Spectroscopy
RARE: Rapid Acquisition with Refocused Echoes
ROI: Region-Of-Interest
RSP: Retrosplenial cortex
SUB: Subiculum
tACS: transcranial Alternating Current Stimulation
tDCS: transcranial Direct Current Stimulation
Thal: Thalamus
TI: Temporal Interference
TI tACS: Temporal Interference transcranial Alternating Current Stimulation
VAPOR: VAriable Power and Optimized Relaxations
VIS: Visual cortex

## Notes

### Competing Interest Statement

The authors have declared no competing interest.

